# Vesicle-templated self-assembly of programmable freestanding multi-µm DNA shells

**DOI:** 10.1101/2025.10.21.683722

**Authors:** Hao Yuan Yang, Christoph Karfusehr, Friedrich C. Simmel

## Abstract

In the quest to create increasingly complex synthetic cell-mimicking systems, a wide range of DNA nanostructures have been developed to coat, permeabilize, sculpt, or otherwise functionalize lipid vesicles. In a complementary strategy, DNA architectures have been used as scaffolds to direct the growth of lipid membrane vesicles. Here we introduce a simple and broadly applicable method to realize freestanding, membrane-mimicking DNA shells: DNA shells are first assembled on the outer surface of giant unilamellar vesicles and then liberated by surfactant-mediated liposome removal. The resulting structures faithfully retain the geometry of their membrane template. We demonstrate the approach with two distinct classes of DNA tectons: a complex barrel-shaped DNA origami with programmable inter-subunit interactions, and a simple nanostar-inspired motif composed of only eleven oligonucleotides. The site-specific addressability of the former enable the rational design of binding interfaces, as demonstrated by controlled multilayer formation. The success of both strategies underscores the generality of our approach and the feasibility of creating shell-like compartments from different DNA architectures. This method enables the construction of tunable, DNA-only containers spanning the size range of eukaryotic cells, offering a fundamentally new type of compartmentalization for bottom-up synthetic biology.

---

Compartmentalization is one of the hallmarks of living systems, where it fulfills a wide range of essential roles. It provides dedicated environments for specific metabolic processes, separates cooperative from parasitic interactions, and prevents the loss of reactants by diffusion, among others. For the same reasons, compartmentalization is also regarded as a prerequisite for the synthetic realization of complex life-like assemblies. Alongside membraneless compartments based on liquid-liquid phase separation, cells typically employ lipid bilayer membranes for compartmentalization. Inspired by this principle, bottom-up synthetic biology has frequently adopted giant unilamellar vesicles (GUVs) as chassis for realizing functional modules that act as synthetic organelles, or even entire synthetic cells [1–4]. However, one of the major challenges for this approach is that loading hydrophilic cargo into the GUV lumen or exchanging materials with the environment requires specialized transport systems, which often limits the implementation of more complex chemical functions [5].

Over the past decade, DNA nanotechnology has provided a versatile toolset for constructing functional modules within synthetic cellular systems. These modules can be used to controllably manipulate membrane architectures and to recapitulate many of the biological functions typically carried out by membrane-associated proteins. The versatility of DNA-based assemblies results from their precisely tunable mechanical properties, high spatial addressability, and rich structural diversity [6–8]. In particular, advances in chemical conjugation strategies [9, 10] have made it possible to attach hydrophobic moieties to defined sites on oligomeric DNA or DNA origami structures, thereby enabling controlled interactions with lipid membranes [11].

DNA origami structures have been interfaced with lipid membranes and vesicles for the formation of artificial nanopores [12–14], to facilitate lipid transfer [15], induce membrane deformation or sculpting [16–18], membrane budding [19], and membrane fusion [20]. Lipid bilayers have been employed to promote the growth of DNA origami lattices by restricting their assembly to two dimensions [21]. DNA nanofilaments have further been used as artificial cytoskeletons, or cell cortexes for synthetic cell models based on vesicles [22–24], droplets [25] or membraneless phase-separated compartments [26, 27]. Conversely, DNA origami structures of various sizes and shapes have been used as scaffolds for the assembly of lipid nanosheets [28] and lipid vesicles of defined sizes [29] or morphologies [30, 31], as well as the production of lipid-bilayer nanodiscs [32].

Previously, liquid-liquid phase-separating DNA nanostars have been used to form a patterned DNA hydrogel layer inside GUVs, which can be released upon surfactant treatment to yield freestanding DNA gel capsules [33]. Such an approach requires the encapsulation of relatively large (*µ*M) DNA concentrations and the use of cationic lipids to promote electrostatic binding complicates applications involving negatively charged cargoes. Furthermore, the resulting capsule walls form disordered, gel-like structures, which makes precise and programmable modification challenging. Once released, the capsules often adopt distorted rather than well-defined spherical shapes. Here we demonstrate an alternative approach in which more complex DNA origami units selectively bind and assemble on the outer surface of lipid bilayer templates, bypassing the need for monomer encapsulation and enabling controlled shell growth prior to release. This strategy yields freestanding multi-*µ*m shells with a well-defined local structure and a preserved, template-guided shape. We first developed a membrane-forming system based on a rigid, precisely addressable membrane-forming DNA origami monomer that can bind to GUV surfaces mediated via cholesterol linkers. This barrel-shaped monomer termed “Dipid” has weak isotropic inter-origami interactions and can mimic lipid membrane assembly, enabling the formation of closed containers reaching the size of *E. coli* cells, multi-*µ*m planar membrane-like sheets, and supports the integration of diverse functional modules [34]. Subsequent removal of the lipid template via surfactant treatment yielded freestanding multi-*µ*m Dipid membrane structures. We also show that the cholesterol-mediated outer-leaflet-templated assembly approach enables the formation of freestanding GUV-sized containers based on a minimal Dipid system composed of only eleven DNA strands per subunit, which robustly retain the shape of their templates.

We defined two criteria for suitable shell-forming monomers: (i) the ability to polymerize into contiguous shells and (ii) the capacity to bind lipid membranes via hybridization with DNA strands conjugated to hydrophobic moieties. These requirements are fulfilled by our Dipid monomers, which are barrel-shaped, hollow structures with a diameter of 30 nm [34, 35]. Dipids assemble into multi-*µ*m monolayers through 30 binding strands extending from the barrel surface [34] (Figure 1). As shown in the figure, each binding strand consists of two distinct subdomains. The sticky domains carry self-complementary sequences that promote binding to other Dipids, largely independent of angular orientation. As the sequences of strands in different layers around the cylindrical body are distinct, they enforce monomer assembly in a single axial orientation. The flex domains, on the other hand, are composed exclusively of thymidines, providing a means to modulate the mechanical properties of the shell. For example, the estimated dimer pitch bending modulus decreases approximately eightfold from *B*_pitch_ = 49 *k_B_T* for 2T flex domains to *B*_pitch_ = 6 *k_B_T* for 16T flex domains (Figure S1). We denote Dipid designs as “PxT”, where *x* refers to the number of T’s in the flex domain, and five different designs were tested (*x* = 2, 4, 8, 16, 32). For incorporation into lipid membranes, Dipid monomers were equipped with three handles protruding from the bottom side of the barrel. These hybridize to linker oligonucleotides carrying a 5’ cholesterol moiety, which are inserted into the GUV membranes (Figure 1a).

**Figure 1.**
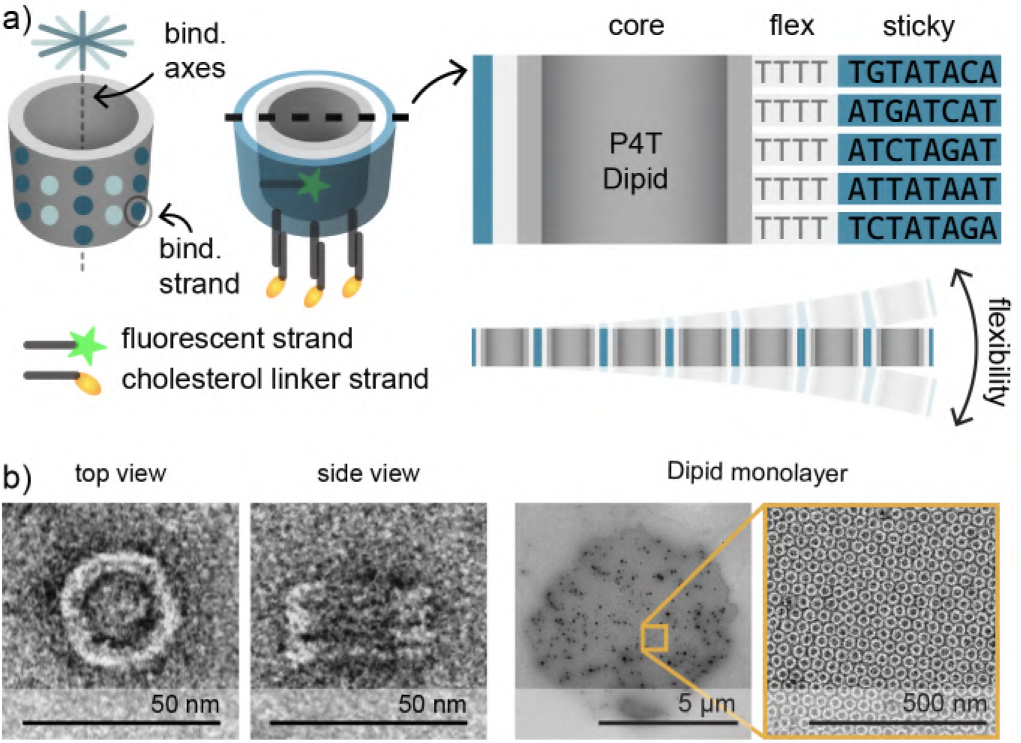
Design of Dipids: programmable and well-structured shell-forming DNA origami monomers. a) Left: Dipids are barrel-shaped, hollow monomers constructed from a core barrel (gray) featuring 30 ssDNA binding strands, distributed on the barrel’s outer surface. Each binding strand consists of a flex domain encoding shell flexibility (light gray) and a sticky domain encoding controlled polymerization (blue). Each Dipid is also functionalized with a fluorophore (green star) conjugated to an internal DNA strand, as well as with handles (dark gray rods) extending from the bottom of the barrel that bind to linker strands conjugated to cholesterol (yellow ellipses). Top right: Cross-section of a P4T Dipid showing the sequences of the flex and sticky domains. Bottom right: A sheet of assembled Dipids schematically showing the flexibility of Dipid membranes. b) Negative stain transmission electron microscopy (TEM) micrographs showing top and side views of the Dipid monomer and a multi-*µ*m assembled Dipid monolayer (P16T) next to a zoom-in.

Non-templated Dipid monomers self-assemble into large 3D aggregates already during the initial folding step. To reassemble Dipid shells on GUV templates, these aggregates must first be disassembled into monomers by heating above a disassembly temperature specific to each Dipid variant. Hence, we performed dynamic light scattering (DLS) measurements for each Dipid variant during both heating and cooling in stepwise increments of Δ*T* = 1 K, with *τ*_1*/*2_ extracted at each step (Figure 2a and S2). During heating, a sudden decrease in the half-decay lag time, *τ*_1*/*2_, indicated disassembly of Dipid aggregates into monomers, whereas during subsequent cooling, a gradual increase in *τ*_1*/*2_ reflected their reassembly.

**Figure 2.**
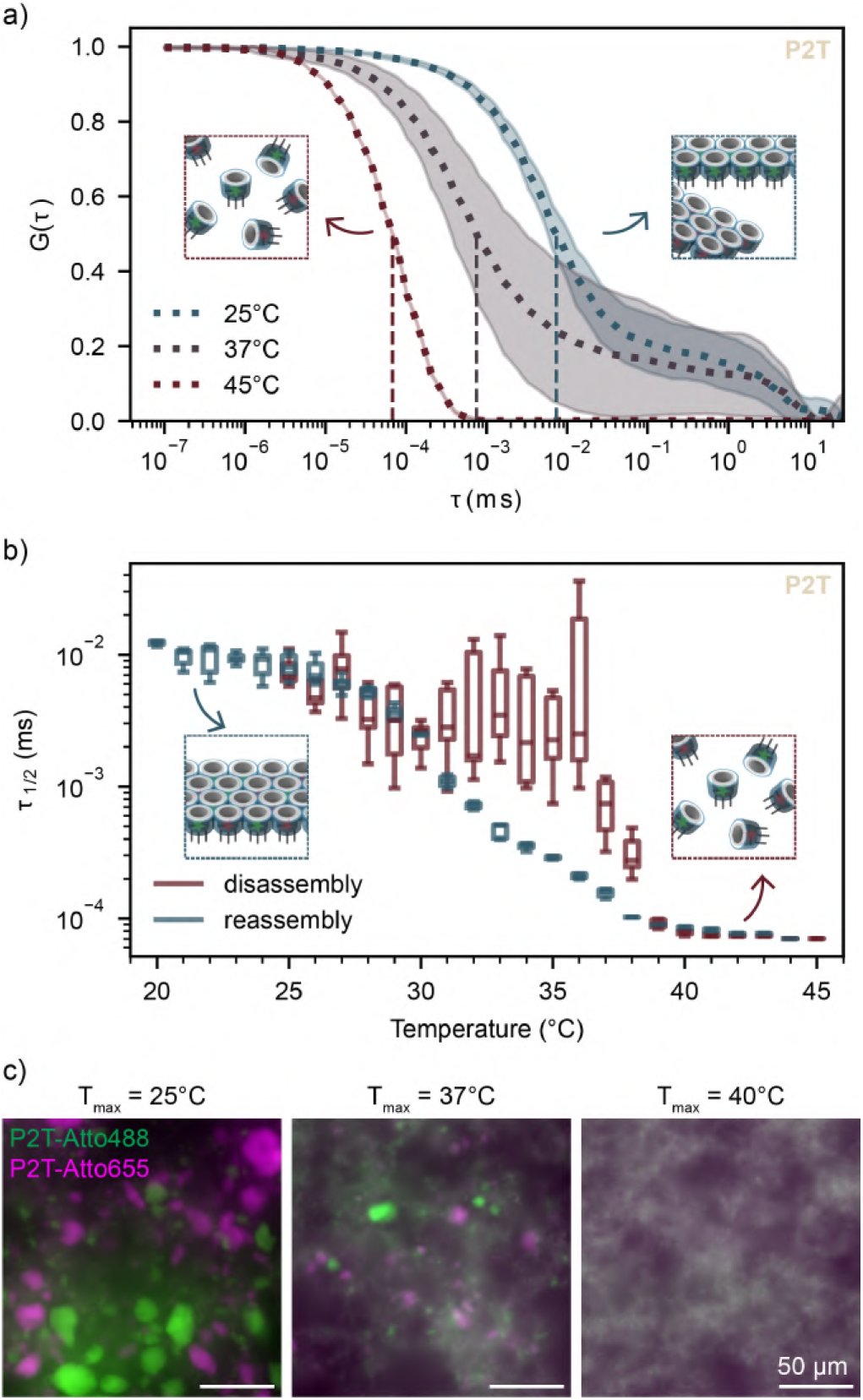
Determination of disassembly temperature. a) Normalized autocorrelation curves of P2T via DLS upon heating up to 25 *^◦^*C (inset showing Dipid aggregates containing different fluorescently-labeled monomers prior to disassembly, 37 *^◦^*C, and 45 *^◦^*C (inset showing disassembled Dipid monomers). Curves represent the statistical mean at any acquisition time, shadows represent the standard deviation of the same data, vertical dotted lines indicate the half-decay lag times, *τ*_1_*_/_*_2_. b) Extracted *τ*_1_*_/_*_2_ of P2T at each temperature step upon disassembly and reassembly via gradual heating (inset showing disassembled Dipid monomers) and cooling (inset showing reassembled Dipid sheets containing a mix of both Dipid monomers). Boxes show medians and interquartile ranges (IQR); whiskers extend to 1.5×IQR. c) Fluorescence microscopy images showing morphology of P2T upon gradual cooling after sample incubation at the respective temperatures *T_max_*. Scale bars: 50 µm.

The experiments indicated nearly complete disassembly of all variants at 40 ^◦^C (Figure 2b and S3). Accord-ingly, 40 ^◦^C was generally used to disassemble Dipid aggregates prior to their reassembly on GUVs. Notably, variants with longer flex domains (P16T and P32T) required lower disassembly temperatures, which may be attributed to the greater entropic contribution of the flexible linkers.

We further confirmed the disassembly and reassembly protocol by fluorescence microscopy. To this end, we mixed two differently labeled samples of the same Dipid variant (labeled with Atto488 or Atto655) and heated them to a defined maximum temperature, *T_max_*. The samples were then gradually cooled to allow liberated monomers to reassemble. Mixed structures containing both labels were expected only if disassembly had occurred during the heating step. As expected, after heating to *T_max_*= 40 ^◦^C all Dipid variants appeared fully mixed, indicating complete disassembly. Heating to intermediate *T_max_*resulted in both mixed structures and smaller chunks of aggregates showing partial disassembly, in agreement with our DLS data (Figure 2c and S4).

Figure 3 illustrates the step-by-step production of freestanding multi-*µ*m DNA shells. To this end, we first separately generated GUVs via electroformation, as well as folded and purified Dipid structures of any variant (P2T is shown as an example). Both GUVs and Dipids were then combined such that Dipid monomers are in excess in a solution composition that prevents unspecific DNA-lipid interactions [17] and reduces GUV sedimentation (Step 1, Figure 3, see also Methods).

**Figure 3.**
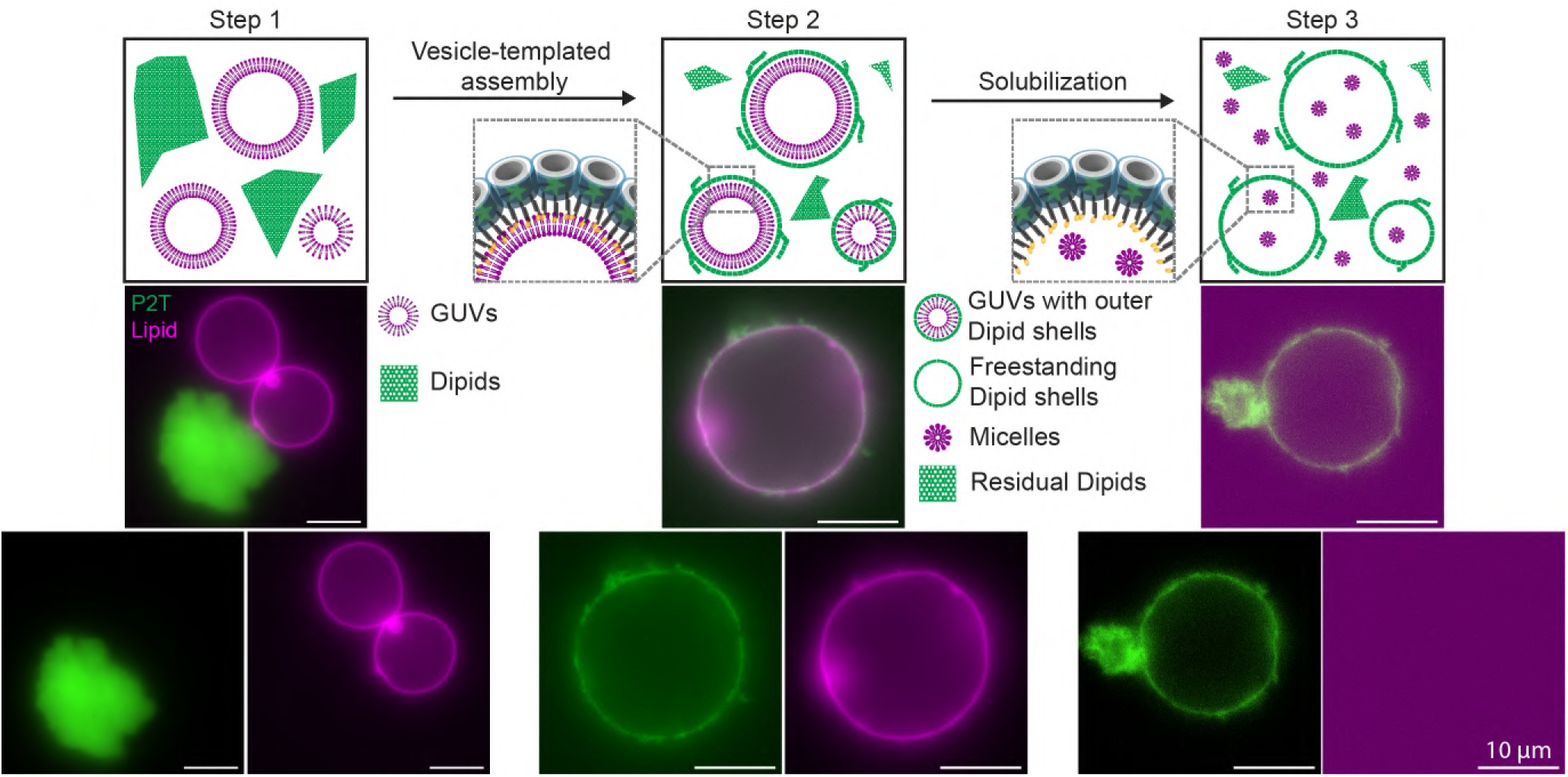
Formation of freestanding DNA shells using the Dipid framework. Schematic illustration of the three-step strategy for the formation of Dipid-based freestanding DNA shells, with accompanying fluorescence images. Step 1: A solution containing GUVs (magenta) and excess Dipids (green) is prepared in a composition where unspecific interaction and GUV sedimentation are reduced (see Methods). Step 2: Heating the solution at 40 *^◦^*C for 1 h induces Dipid disassembly, and gradual cooling by *−*0.1 *^◦^*C every 3 min facilitates vesicle-templated assembly of Dipids, forming GUVs enclosed by an outer Dipid shell. Step 3: Upon surfactant treatment using 0.1 % Triton X–100, GUVs are solubilized into micelles, resulting in freestanding multi-*µ*m DNA shells. Scale bars: 10 µm.

Subsequently, the Dipid structures were disassembled by heating to 40 ^◦^C. During gradual cooling, the monomers reassembled on the surface of GUVs and GUV aggregates, facilitated by hybridization of their handles with membrane-anchored cholesterol linkers, thereby forming an outer Dipid shell (Step 2, Figure 3 and S5). The fuzzy appearance of the shell likely reflects Dipid monolayer outgrowths, previously observed in self-assembled Dipid membrane clusters [34]. These outgrowths are seeded by origami dimers or other misfolded structures and could, in principle, be suppressed by improved monomer purification.

Following solubilization of the GUVs into micelles by addition of the surfactant Triton X–100, we obtained freestanding multi-*µ*m DNA shells without any detectable lipid signal at the shell (Step 3, Figure 3 and S6). The resulting shells appeared contiguous (Figure S7). However, during solubilization, some DNA shells collapsed. As the Dipids were linked to the GUVs via cholesterol-binding handles, the forces generated in the release process likely caused this collapse, particularly in the case of P32T, possibly due to its higher structural flexibility (Figure S6).

We additionally demonstrated that each component of the Dipid monomer design is required for correct assembly: the absence of cholesterol-binding handles or leaving the reaction at room temperature without first inducing Dipid disassembly prevented shell formation, whereas the absence of sticky domains in the monomer (“P8Td”) did not result in the formation of freestanding Dipid shells after solubilization (Figure S8).

Cryo-ET structural studies have shown that Dipids indeed self-assemble into monolayers as designed [34]. We therefore surmised that templated growth of Dipids would also occur in monolayers. To test this hypothesis, we compared the fluorescence signal of labeled Dipids on GUV templates with that of monolayers formed by non-polymerizing P8Td Dipids (Figure 4a,b). Compared to the monolayer control, the median Dipid shell intensity is *≈* 1.2 times higher and exhibits a broader distribution(I_P8T_ = 3547 *±* 1689 a.u., I_P8Td_ = 3026 *±* 950 a.u.). This likely reflects additional intensity contributions from membrane outgrowths, suggesting that otherwise the Dipids are organized in monolayers.

**Figure 4.**
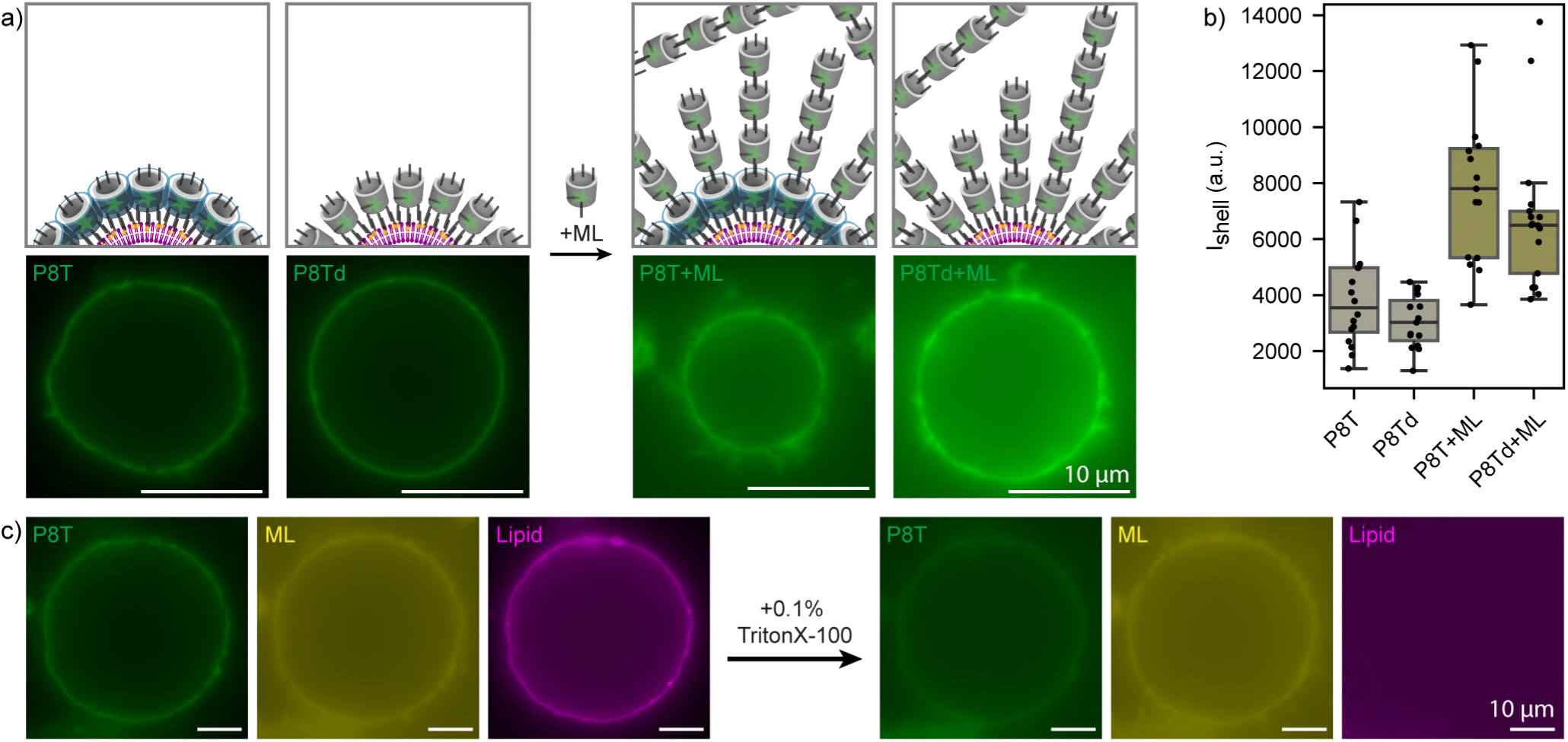
Formation and release of multilayer shells. a) Schematics showing comparison of P8T Dipid shells (green) with non-polymerizing P8Td Dipid shells (green) assembled on GUVs before and after addition of a multilayer Dipid variant (“ML”, green) capable of radial polymerization. Corresponding fluorescence microscopy images were recorded under identical conditions and are shown with the same contrast settings, with a representative image shown. b) Median background-corrected fluorescence intensities of Dipid layers of analyzed templated structures. Boxes show medians and interquartile ranges (IQR); whiskers span 1.5×IQR. Sample sizes: n_P8T_ = 16, n_P8Td_ = 15, n_P8T+ML_ = 15, and n_P8Td+ML_ = 17. Comparison of the shell-forming P8T and non-polymerizing P8Td structures suggests that the P8T Dipids indeed assembled into a monolayer. c) Fluorescence microscopy images of a P8T DNA shell (green) assembled on a GUV (magenta) with a multilayer Dipid shell on the outside (yellow) before and after release using Triton X–100. Scale bars: 10 µm.

By exploiting the site-specific addressability of DNA origami structures, the first monolayer can serve as a starting point for the realization of more complex multilayer shells. Notably, exposure of the first layer to the surrounding solution allows easy access for implementing layer-specific modifications. As a proof of principle, we developed a multilayer (“ML”) Dipid that binds to the top of shell Dipids and polymerizes top-to-bottom with copies of itself (Figure 4a and Figure S9). The addition of ML Dipids to preformed Dipid shells increased shell thickness, evident from increased shell intensities (Figure 4a,b, I_P8T+ML_ = 2.2 × I_P8T_, I_P8Td+ML_ = 1.8 × I_P8Td_). Releasing such multilayer structures produced our largest freestanding Dipid shells (diameter *≈*40 µm) (Figure 4c and Figure S10, S11), which notably retained the size and shape of their GUV templates (Figure 4c). If finer layer control is desired, the Dipid shell system enables stepwise addition of individual Dipid layers carrying small molecules such as fluorescent dyes (cf. Figure S9, S10, for a two-layer system) or other functional units.

We hypothesized that membrane-like structures could also be formed by other types of nucleic acid assemblies, not necessarily restricted to our approximately 140-component DNA origami Dipid design. To test whether our assembly method is applicable to less structured systems, we designed a minimal monomer inspired by DNA nanostars (Figure 5a and S12). This monomer consists of 11 DNA strands and assembles into a miniature Dipid with six binding domains and additional ssDNA loop that hybridizes to a cholesterol-modified DNA strand. We simulated the structure of the minimal monomer using oxDNA [36], which suggests that the designed structure can form experimentally.

**Figure 5.**
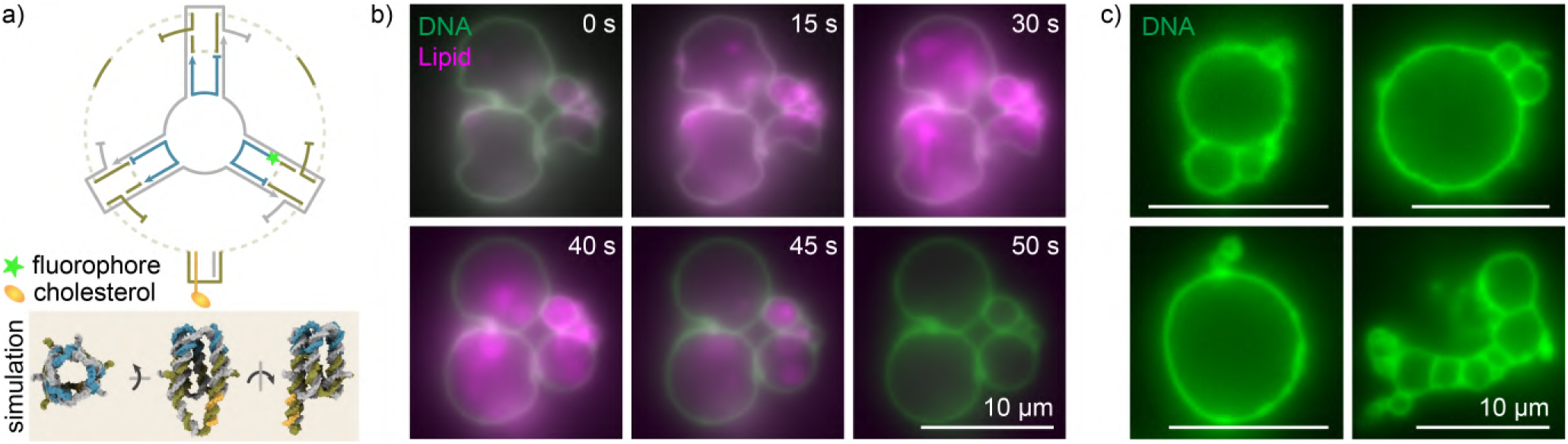
Formation of freestanding DNA shells from a minimal monomer. a) DNA strand routing of the minimal monomer and mean structure from an oxDNA simulation. Strands are colored to highlight routing symmetries. Dotted lines represent strand domain connections of zero length. Binding domains have the sequence “TT GCGC” (Figure S9). b) Fluorescence time series of DNA shell (green) release from GUV templates (magenta). Triton X–100 was added to one side of a narrow channel containing GUVs enclosed by DNA shells and allowed to diffuse. *t* = 0 s marks the start of acquisition. c) Examples of freestanding DNA shells after Triton X–100 treatment. Scale bars: 10 µm.

We reasoned that the formation of a contiguous DNA shell does not strongly depend on the details of the monomer structure, provided that the monomers can assemble into extended sheets with sufficient thickness and flexibility. We expected that our miniature Dipids would yield such sheets through their symmetric and approximately isotropic inter-subunit interactions, which are realized by their six self-complementary binding domains. To test this, we implemented a one-pot protocol that combined monomer folding with vesicle-templated shell assembly in a single annealing step. As expected, this yielded DNA shells assembled on both GUVs and GUV aggregates (Figure 5b).

Upon treatment with surfactant, the GUV templates dissociated from the DNA shells, passing through intermediates that included tubular structures and smaller liposomes, and ultimately leaving freestanding DNA shells (Figure 5b,c) with no detectable lipid signal at the DNA boundary (Figure S13a,b). Notably, we sometimes observed non-spherical templated structures that relaxed into spherical shells after liposome destruction, thus retaining the sizes and shapes of the GUV templates (Figure 5b).

In summary, we have developed a straightforward and versatile strategy to create freestanding, eukaryotic cell-sized DNA shells by assembling DNA nanostructures on the outer surfaces of lipid vesicles, followed by their release via surfactant treatment. Templated self-assembly on the outer GUV leaflet, mediated by hydrophobic linkers, enforces a preferred monomer orientation and enables polymerization under buffer conditions that suppress unspecific DNA-lipid interactions. Together with programmable Dipid monomer units, this produces membrane-like DNA shells with a hexagonal lattice organization. The defined orientation and precise geometry of Dipids and Dipid membranes support the rational design of binding interfaces, in principle allowing intra-Dipid-membrane integration of functional modules [34]. In addition, accessibility from the outside enables the sequential addition of functionalized shell layers.

Unlike nanostar-based gel capsules, our DNA shells retain the shapes and sizes of their lipid templates upon release. We observed distinct mechanical responses for the two monomer types used (Dipid and the minimal monomer). While both types of DNA shells withstand moderate deformation, shells formed from the minimal monomer accommodate stronger template deformations and relax back to a spherical geometry after lipid removal. We therefore anticipate that monomer design will enable programmable shell properties - such as membrane stiffness, porosity, and geometry retention - tailored to specific applications.

The system may be further advanced by mitigating outgrowth formation, for instance by more extensive purification of Dipid monomers or by applying temperature-cycling and washing steps to remove unbound species. Although such outgrowths could, in principle, be exploited as high-surface-area features, they may interfere with downstream shell processing or with the integration of functional modules. In our experiments, designed multilayer growth considerably improved mechanical robustness and reduced shell collapse. We anticipate that multilayer assembly may be further enhanced by designing different types of layer-specific Dipids and supplying them to immobilized GUV templates in a microfluidic device to enable layer-by-layer assembly. Moreover, photocleavable or pH-responsive cholesterol linkers offer alternative release strategies that may further reduce collapse during lipid removal.

As our templating strategy only requires an accessible lipid bilayer, we anticipate that it may also be applicable to natural cells or organelles with complex geometries. By adding functional Dipid modules and applying the Dipid organization principles in binary mixtures developed in our previous work [34], we foresee that cell-sized functional DNA compartments could be realized in the near future. In the longer term, composite DNA shells incorporating biological and synthetic components may provide a route toward bio-hybrid compartments that support increasingly complex, programmable behaviors and could contribute to the development of biomolecular robotic systems.

## Acknowledgments

F.C.S. gratefully acknowledges support by and H.Y.Y. and C.K. conducted their research within the Max Planck School Matter to Life supported by the German Federal Ministry of Research, Technology and Space (BMFTR) in collaboration with the Max Planck Society.

## Data availability

All data are available from the authors upon request.

## Supplementary Information

### Materials and Methods

#### oxDNA simulations

The nanostructure of the minimal monomer was visualized with oxView [37], which was also used to define the relaxation forces used during the subsequent oxDNA simulation. Following an initial Monte Carlo pre-relaxation, we performed molecular dynamics simulations using the oxDNA2 model [36] to relax and simulate the monomer structure. We generated PDB files for visualization in ChimeraX [38] via the oxDNA analysis tools command-line script [36]. The shown monomer structure corresponds to a mean configuration. All relevant oxDNA files are available on our GitHub repository.

#### DNA origami design and folding

We incorporated the modifications mentioned in the main text into the Dipid structures [34], which were in turn adapted from the DNA origami structures developed by Wickham et al. [35] using scadnano [39]. Dipids were folded with a p2873 (30 nm Dipid) scaffold (provided by Prof. Hendrik Dietz’s group, 100 nM in ddH_2_O) and varying sets of staple strand oligonucleotides (Integrated DNA Technologies, 200 µM in 20 mM TRIS, 0.1 mM EDTA, pH 8.0). Annotated scadnano files and all DNA sequences used in this study are available on our GitHub repository. We prepared 60 µL folding solutions containing a final scaffold concentration of 50 nM and a staple strand concentration of 200 nM in FOB18 buffer (5 mM TRIS, 1 mM EDTA, 18 mM MgCl_2_, and 5 mM NaCl). Folding solutions were annealed in a thermocycler (Mastercycler nexus GX2, Eppendorf) using a protocol of 15 min at 65 ^◦^C followed by a decrease of 0.1 ^◦^C every 6 min from 56 ^◦^C to 53 ^◦^C.

#### DNA origami purification via ultrafiltration

Folded Dipid samples were purified via ultrafiltration using Amicon Ultra filters (0.5 mL, 100 K, Millipore) in 32 ^◦^C FOB5 washing buffer (5 mM TRIS, 1 mM EDTA, 5 mM MgCl_2_, 5 mM NaCl). The filters were pre-washed using 500 µL FOB5, then loaded with 430 µL of FOB5 and 60 µL of folding solution, followed by 450 µL of FOB5, and finally 450 µL of FOB5_NaCl300 (5 mM TRIS, 1 mM EDTA, 5 mM MgCl_2_, 300 mM NaCl), with centrifugation at 20 krcf for 5 min at 32 ^◦^C and discarding of flowthrough done at each step. The purified and buffer-exchanged samples were then extracted from the filters by repeated aspiration to dissolve any pellets.

#### DNA origami purification via Polyethylene Glycol (PEG) precipitation

Alternatively, we purified folded Dipid samples via PEG precipitation. Up to four folding reactions of the same Dipid type and FOB18 buffer were combined up to a volume of 750 µL. This was mixed with 750 µL of precipitation buffer (15 % *w/v* PEG, 500 mM NaCl, 1x FOB0), and centrifuged at 20 krcf for 30 min at 25 ^◦^C. After removing the supernatant, and the pellet was resuspended in 750 µL of FOB18 and incubated at 30 ^◦^C, 500 rpm for 30 min. Afterwards, another 750 µL of precipitation buffer was added, and the centrifugation step was repeated. The supernatant was discarded, and the sample was resuspended in at least 50 µL of FOB5_NaCl300 by incubating at 30 ^◦^C, 500 rpm for a minimum of 30 min.

#### Negative stain TEM

We applied 5 µL of DNA origami solution to glow-discharged formvar carbon Cu400 TEM grids (Science Services) with a coating time of 20 s, a coating current of 35 mA, and negative polarity. The incubation time ranged from 30 to 300 s, depending on the DNA origami concentration. To prepare the staining solution, we added 1 µL of 5 M NaOH to 200 µL of 2 % uranyl formate, then vortexed and centrifuged the mixture at 21 krcf for 5 min. After sample incubation, we washed the grids with 5 µL of stain and subsequently incubated them with 15 µL of stain for 30 s. nsTEM imaging was conducted using a FEI Tecnai T12 microscope (120 kV) equipped with a Tietz TEMCAM-F416 camera and operated via SerialEM.

#### Dynamic light scattering (DLS)

We performed DLS measurements using a DynaPro NanoStar (Wyatt Technology), using the internal temperature control to disassemble and reassemble Dipid solutions (7.5 nM in 80 µL of FOB5_NaCl300). Dipid solutions were added to disposable MicroCuvettes (Wyatt Technology), with the chamber sealed to prevent evaporation. The same protocol was used for all Dipid variants: a slow temperature ramp of 1 ^◦^C per 10 min from 25 ^◦^C to 45 ^◦^C to 20 ^◦^C, with acquisition starting after 5 min at each step to allow the target temperature to be reached. At each temperature step, we acquired 10 acquisitions with a 30 s acquisition time and with auto-attenuation enabled. We extracted raw DLS autocorrelation curves from the instrument files, applied moving average smoothing, and performed min–max normalization of the smoothed data. For each measurement, we determined the autocorrelation time *τ*_1*/*2_, defined as the time at which *G*(*τ*) *≤* 0.5 *G*(*τ* = 0).

#### GUV formation by electroformation

For all Dipid experiments, we produced GUVs by electroformation. We prepared a stock solution of 0.988% DOPC (1,2-dioleoyl-sn-glycero-3-phosphocholine, Avanti Research), 0.01% Atto655-labeled DOPE (1,2-dioleoyl-sn-glycero-3-phosphoethanolamine, ATTO-TEC), 0.002% Chol-TEG linker (oligonucleotide carrying a 5’ cholesterol moiety connected via a tetraethylene glycol spacer, Biomers) in chloroform at a total concentration of 2 mM. We spread 5 µL of the solution evenly on platinum electrodes before drying in a desiccator. The electrodes were then immersed in 650 µL of 660 mM sucrose, before applying 20 Vpp at 10 Hz for 60-90 min followed by 1 Hz for another 30 min at room temperature using a custom electroformation chamber and a function generator (FG-1302, Voltcraft). We stored GUV solutions at 4 ^◦^C till further use.

#### GUV formation by natural swelling

For all minimal monomer experiments, we prepared GUVs by natural swelling. Specifically, we added 150 µL of 2 mM POPC (1-palmitoyl-2-oleoyl-glycero-3-phosphocholine, Avanti Research) in chloroform containing 0.5 % Atto488-labeled DOPE (ATTO-TEC) to a small round-bottom flask. We dried the lipid film using a rotary evaporator at 100 mbar for 3 h. To obtain a final lipid concentration of 100 µM, we hydrated the film with 3 mL 1 M sucrose and incubated it at 4^◦^ overnight.

#### Dipid shell formation

We combined 50 µL reaction solutions in a final concentration of 1.5 nM Dipids, 10% Optiprep (Serumwerk), 40 % (v/v) GUVs containing TEG-chol linkers produced via electroformation in 660 mM sucrose (final concentration of 264 mM sucrose), in 1× FOB5_NaCl300. This composition matches the osmolarity of the inner solution of the GUVs (660 mM sucrose), as determined by an osmometer (Osmomat 3000, GONOTEC GmbH), while Optiprep helps to reduce sedimentation of GUVs. The presence of 300 mM NaCl serves to prevent unspecific DNA-lipid interactions [17]. We then performed vesicle-templated assembly by incubating reaction solutions in a thermocycler using a protocol of 1 h at 40 ^◦^C followed by a decrease of 1 ^◦^C every 3 min from to 20 ^◦^C, forming GUVs with outer Dipid shells. For fluorescence imaging, 20 µL of the annealed reaction solution was added to 50 µL of imaging buffer (5 mM TRIS, 1 mM EDTA, 5 mM MgCl_2_, 300 mM NaCl, 264 mM sucrose) into microscopy slide wells (ibiTreat µ-slide 18 well, Ibidi).

#### Release of Dipid shells

To release freestanding Dipid shells, we gently added 0.7 µL of Triton X–100 (1:9 diluted with FOB5_NaCl300, final 0.1%, Sigma-Aldrich) to the above-mentioned 70 µL of imaging solutions containing GUVs with outer Dipid shells.

#### Bilayer and multilayer Dipid shell formation and release

To form bilayer or multilayer Dipid shells, we mixed 5× excess of 1L Dipids or 25× excess of ML Dipids (pre-heated to 50 ^◦^C) respectively with solutions containing P8T Dipids assembled on GUVs in a final volume of 100 µL with the same buffer composition. The mixtures were left at room temperature for at least two hours before the addition of 100 µL imaging buffer in microscopy slide wells (ibiTreat µ-slide 18 well, Ibidi) for fluorescence imaging. To release freestanding bilayer or multilayer Dipid shells, we similarly added 2 µL of Triton X–100 (1:9 diluted with FOB5_NaCl300, final 0.1%, Sigma-Aldrich) to the above-mentioned 200 µL of imaging solutions.

#### Minimal monomer shell formation

We combined 2 µL of 100 µM cholesterol strand solution (Biomers, resuspended in FOB0_NaCl500; 5 mM TRIS, 1 mM EDTA, 500 mM NaCl) with 18 µL GUV solution produced using natural swelling, and incubated the mixture for 20 min at room temperature. The one-pot folding and shell assembly mixture contained the DNA oligonucleotides (Integrated DNA Technologies, 200 µM in 20 mM TRIS, 0.1 mM EDTA, pH 8.0) “sticky1” to “sticky6”, “dummy”, “staple1”, and “staple2” at 0.888 µM, “staple3_fluo” at 1 µM, 1× FOB_NaCl500, and 16.67 % (v/v) of the GUV solution containing cholesterol strands. We annealed the mixture in a thermocycler from 85 ^◦^C to 25 ^◦^C at *−*1 ^◦^C h^−1^.

#### Release of minimal monomer shells

The solution containing GUVs enclosed by minimal monomer shells was transferred to channels of a microscopy slide (µ-Slide VI 0.1, Ibidi). We added 5 µL Triton X–100 (1:100 diluted with FOB0_NaCl500) to one side of the channel and allowed it to diffuse into the channel. We recorded time series in regions where the lipid signal and GUV appearance were unchanged and waited for the surfactant to reach these positions and solubilize the GUVs. We also recorded images of freestanding DNA shells in regions with high Triton X–100 concentration and no discernible lipid vesicle signal.

#### Fluorescence microscopy

Each DNA monomer was designed to include one Atto488, Atto565, or Atto655-labeled DNA oligonucleotide (Biomers), while GUVs also contain either Atto488 or Atto655-labeled DOPE (ATTO-TEC). We imaged samples using a Nikon Ti-2E inverted fluorescence microscope (NIS elements software, SOLA SM II LED light source, Andor NEO 5.5 camera) with a 40× air objective (CFI Plan Fluo 40× CH, Nikon) or a 100× oil objective (Lambda D 100× Oil CFI Plan Apochrom, Nikon). Images were globally contrast-adjusted using Fiji [40].

#### Validation of Dipid disassembly temperatures via fluorescence microscopy

For imaging of Dipid structures after disassembly and reassembly to verify disassembly temperatures, we separately folded and purified two differently-labeled (Atto488 or Atto655) samples of the same Dipid variant, then combined them in approximately equal ratios in a final concentration of 7.5 nM in FOB5_NaCl300. The solutions were then incubated at the specified temperatures for 1 h, followed by cooling at *−*0.1 ^◦^C every 3 min to 20 ^◦^C in a thermocycler. Solutions were transferred into microscopy slide wells (ibiTreat µ-slide 18 well, Ibidi) prior to fluorescence microscopy imaging.

---

**Figure S1.**
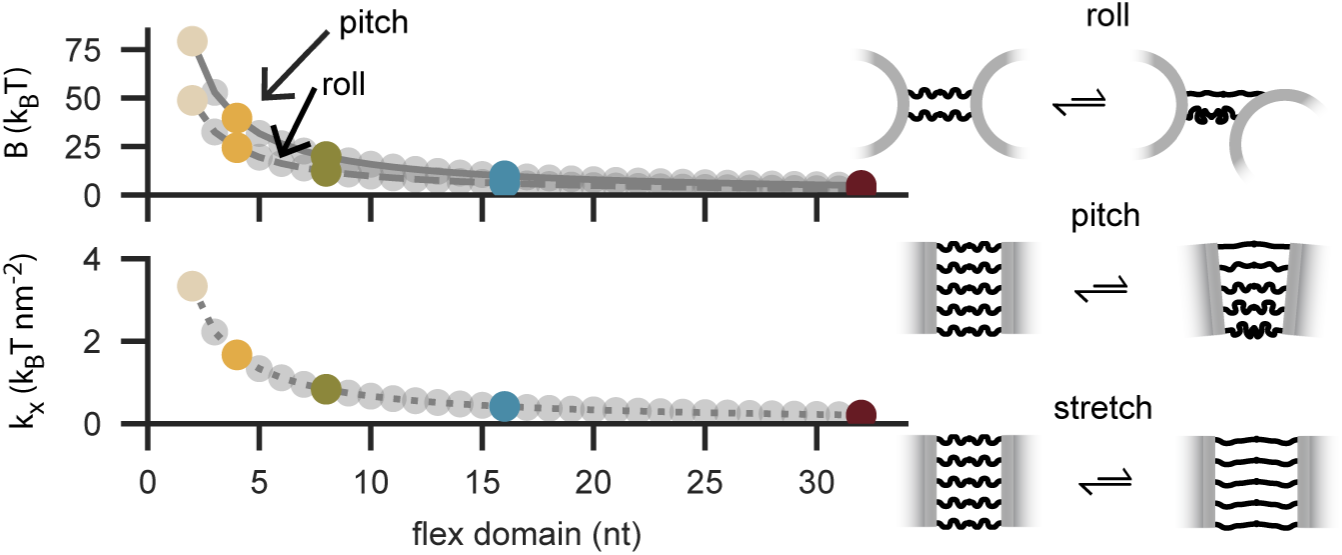
Estimates of selected mechanical properties of Dipid dimers. Schematics of the two bending modes and the stretching mode used to estimate the bending modulus *B* for each mode and the stretching modulus *k_x_*, respectively. PxT Dipid designs used in this study are marked by colored circles. All estimates follow the framework of Videbæk *et al.* [41], adapted to Dipid geometry.

**Figure S2.**
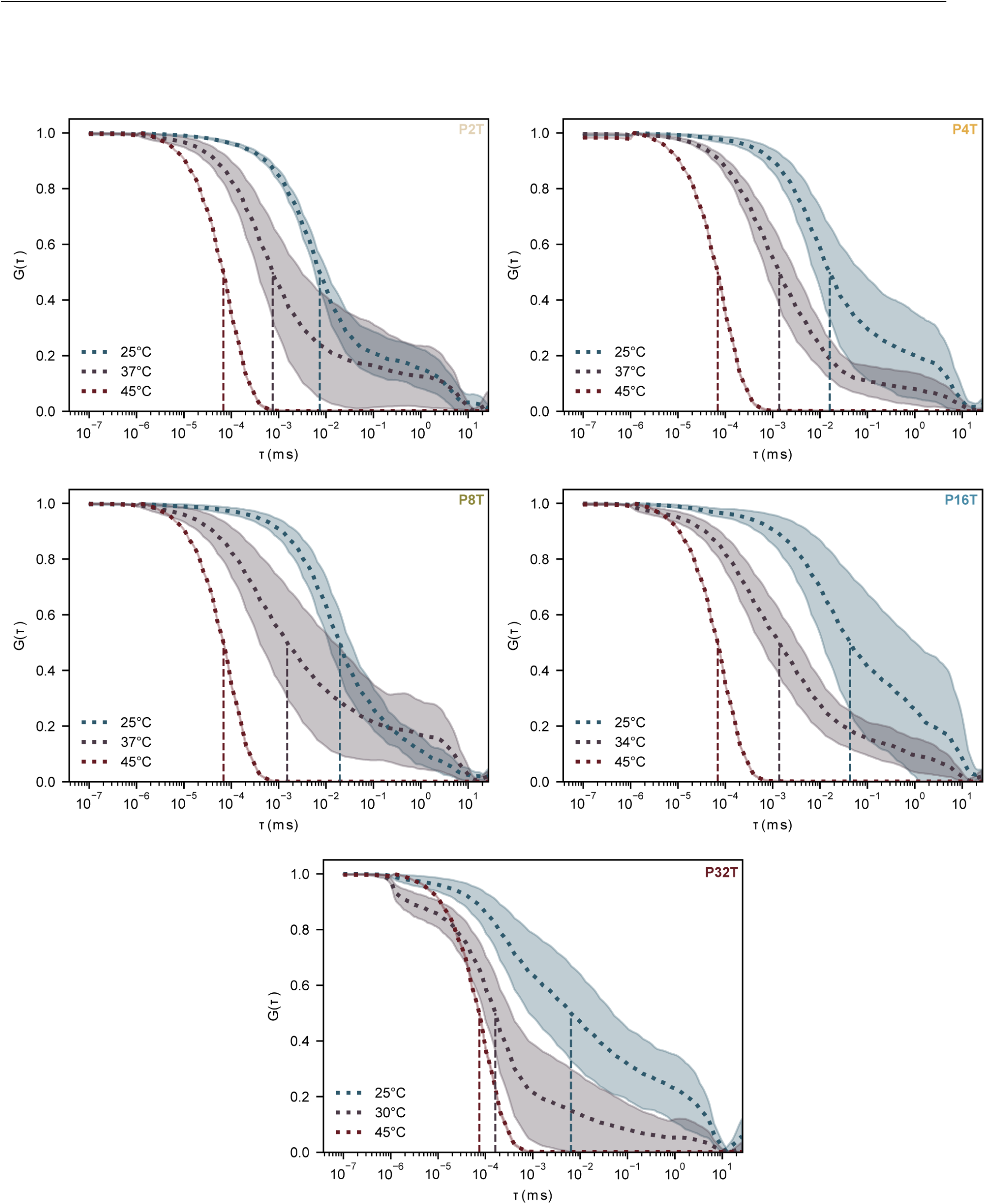
Normalized autocorrelation curves obtained via DLS during the disassembly of each Dipid variant at 25 *^◦^*C, an intermediate temperature, and 45 *^◦^*C. Curves represent the statistical mean at any acquisition time, shadows represent the standard deviation of the same data, vertical dotted lines indicate the half-decay lag times, *τ*_1_*_/_*_2_.

**Figure S3.**
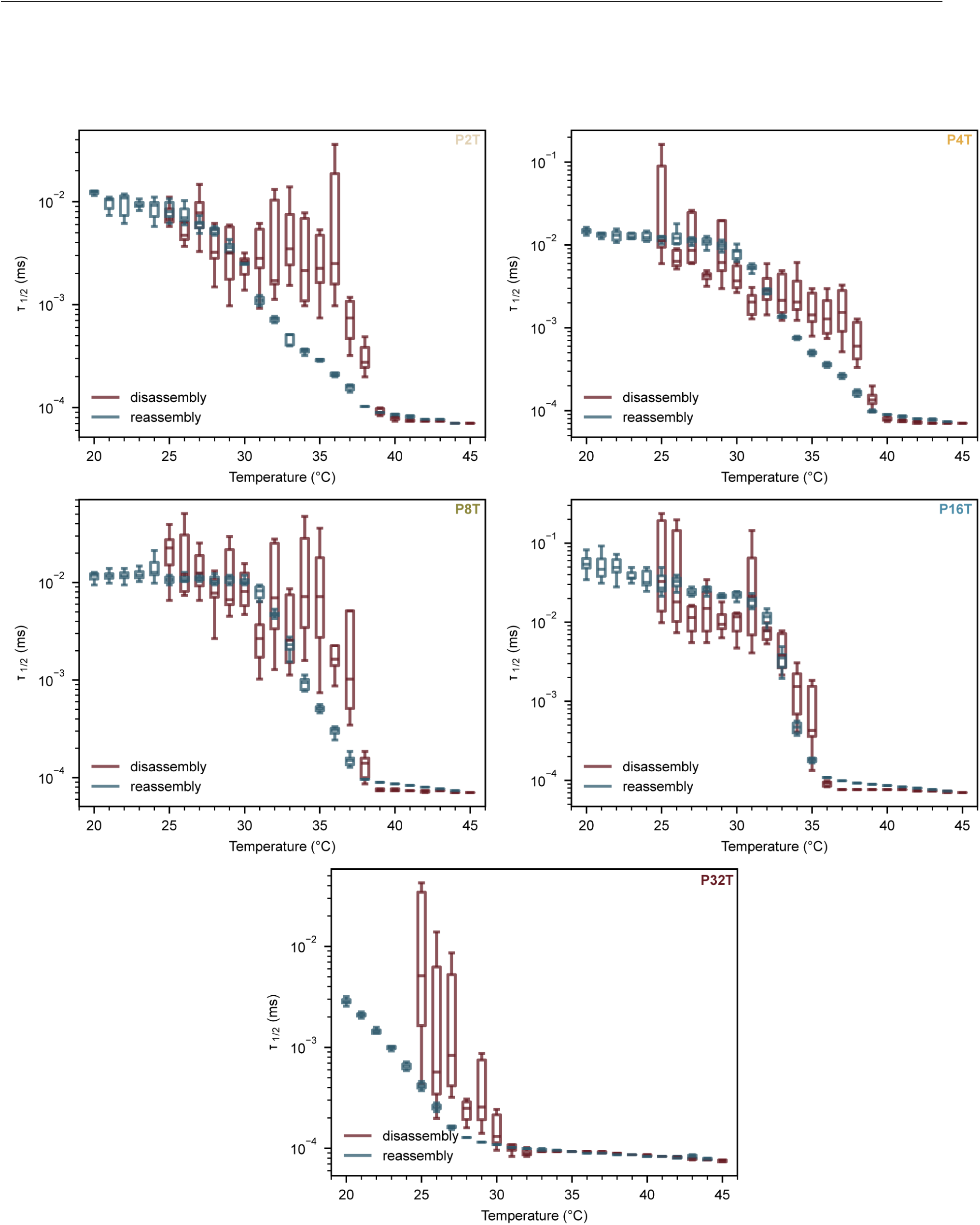
Extracted half-decay lag times, *τ*_1_*_/_*_2_, of each Dipid variant upon disassembly and reassembly at each temperature step obtained via DLS. Boxes show medians and interquartile ranges (IQR), whiskers extend to 1.5×IQR.

**Figure S4.**
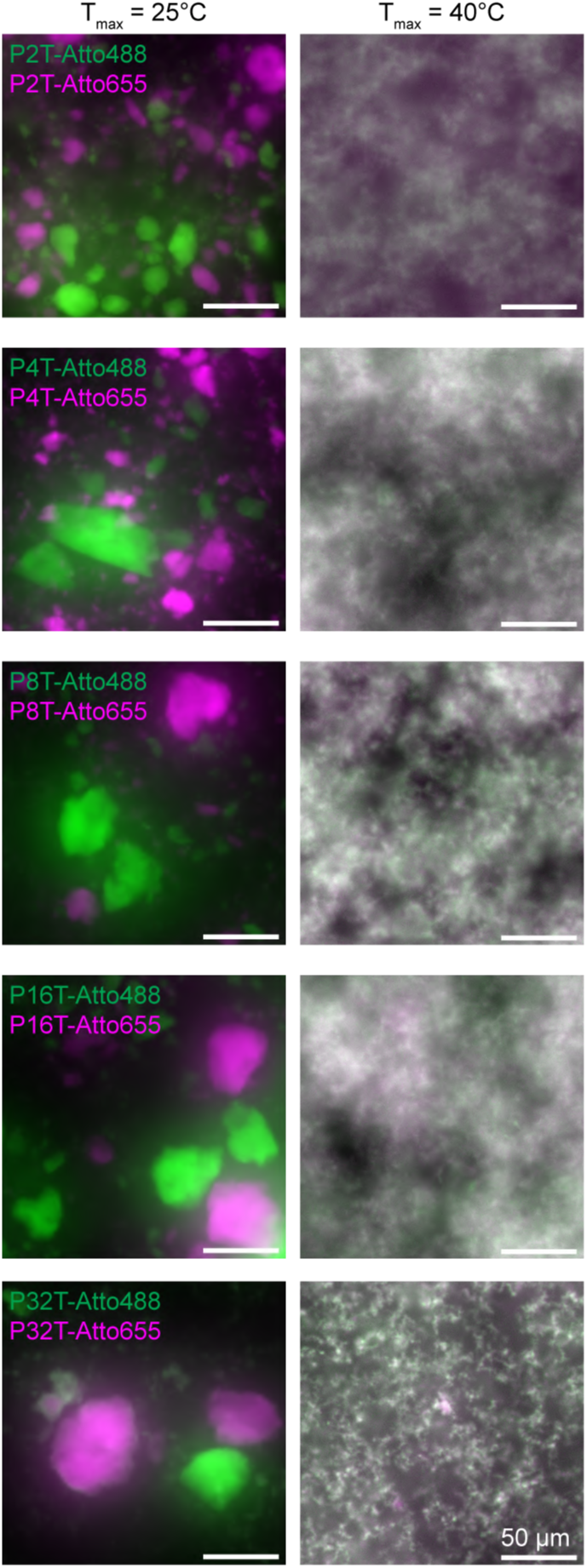
Fluorescence microscopy images of each Dipid variant, each containing a mix of structures labeled with Atto488 (green) or Atto655 (magenta), upon gradual cooling after incubation at the indicated temperatures *T_max_* for 1 h. Scale bars: 50 µm.

**Figure S5.**
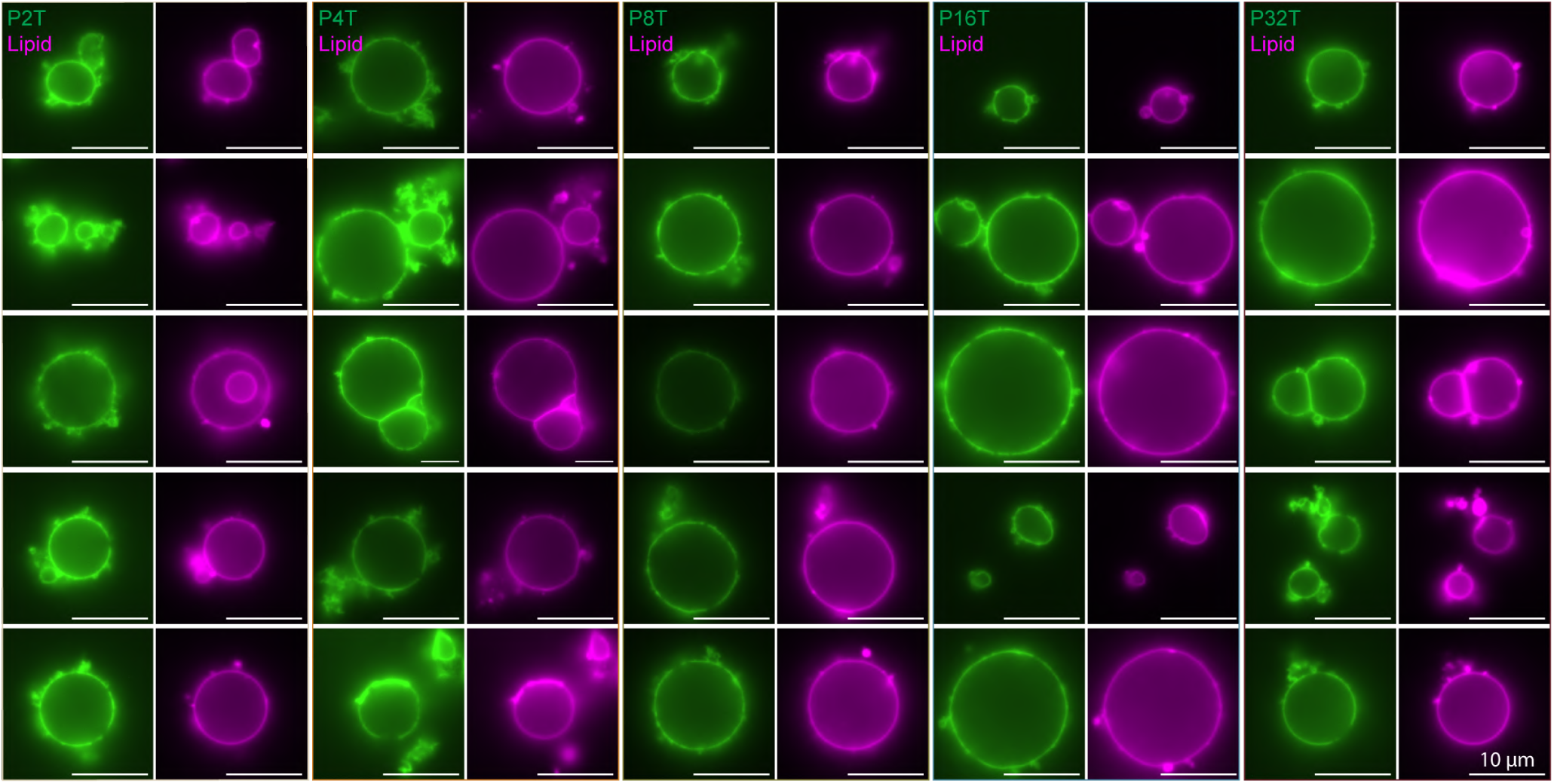
Fluorescence microscopy images of each Dipid variant (green), upon reassembly on GUVs (magenta), forming outer DNA shells. Scale bars: 10 µm.

**Figure S6.**
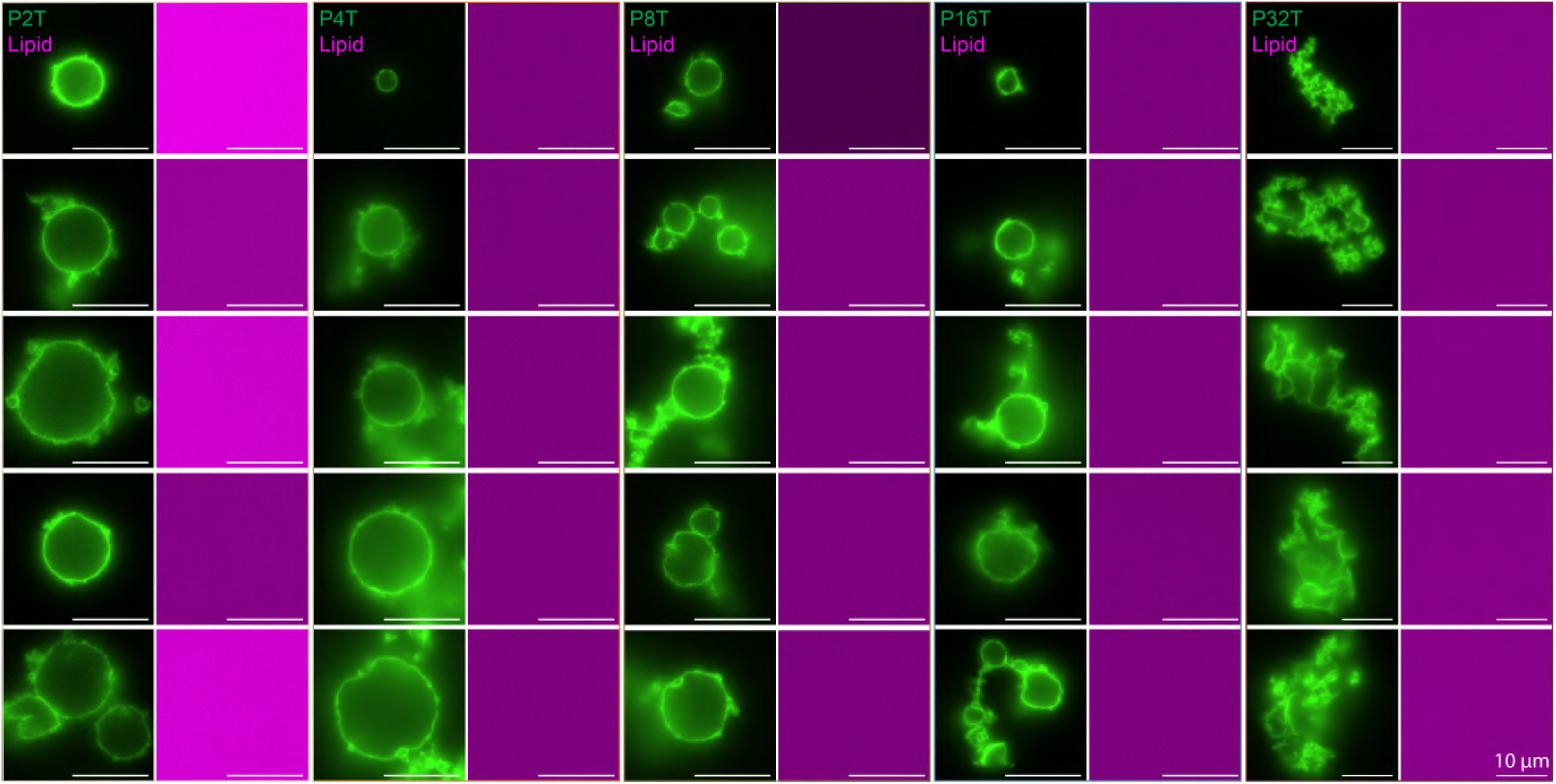
Fluorescence microscopy images of each Dipid variant (green), upon solubilization of GUVs (magenta) after vesicle-templated assembly, forming freestanding spherical Dipid shells with the exception of P32T, where Dipid shells collapsed. Scale bars: 10 µm.

**Figure S7.**
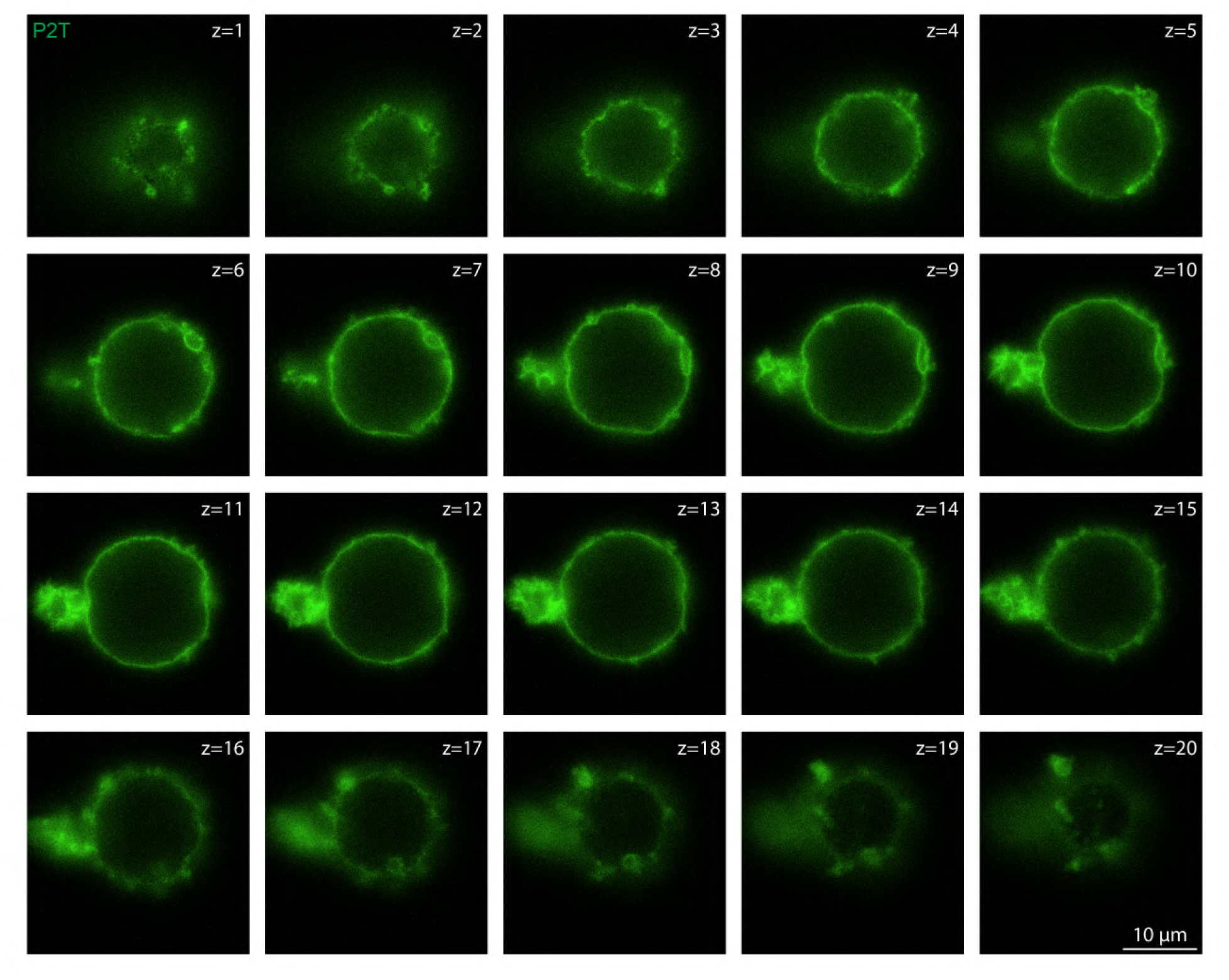
Fluorescence microscopy Z-slices of the freestanding Dipid shell (green) from Figure 3 (Δ*z* = 1 µm). Scale bar: 10 µm.

**Figure S8.**
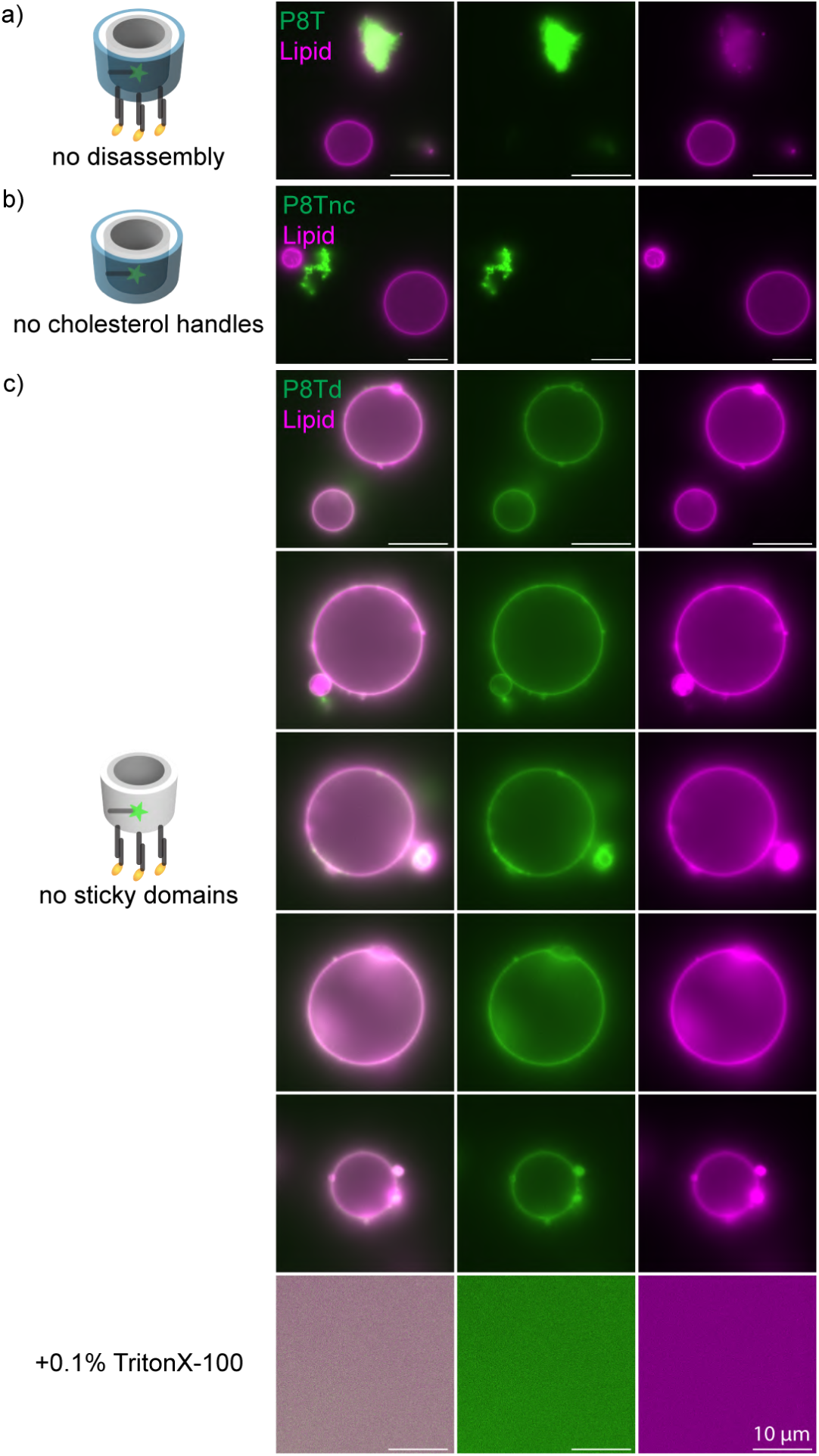
Fluorescence microscopy images showing no formation of Dipid shells (green) enclosing GUVs (magenta) when a) no disassembly occurs for P8T, or when b) cholesterol handles (dark gray rods) protruding from the bottom of the Dipid barrel are absent (“P8Tnc”), as well as no freestanding Dipid shells after Triton X–100 addition when c) sticky domains (blue cylinder layer) are absent (“P8Td”) despite formation of outer Dipid shells. Scale bars: 10 µm.

**Figure S9.**
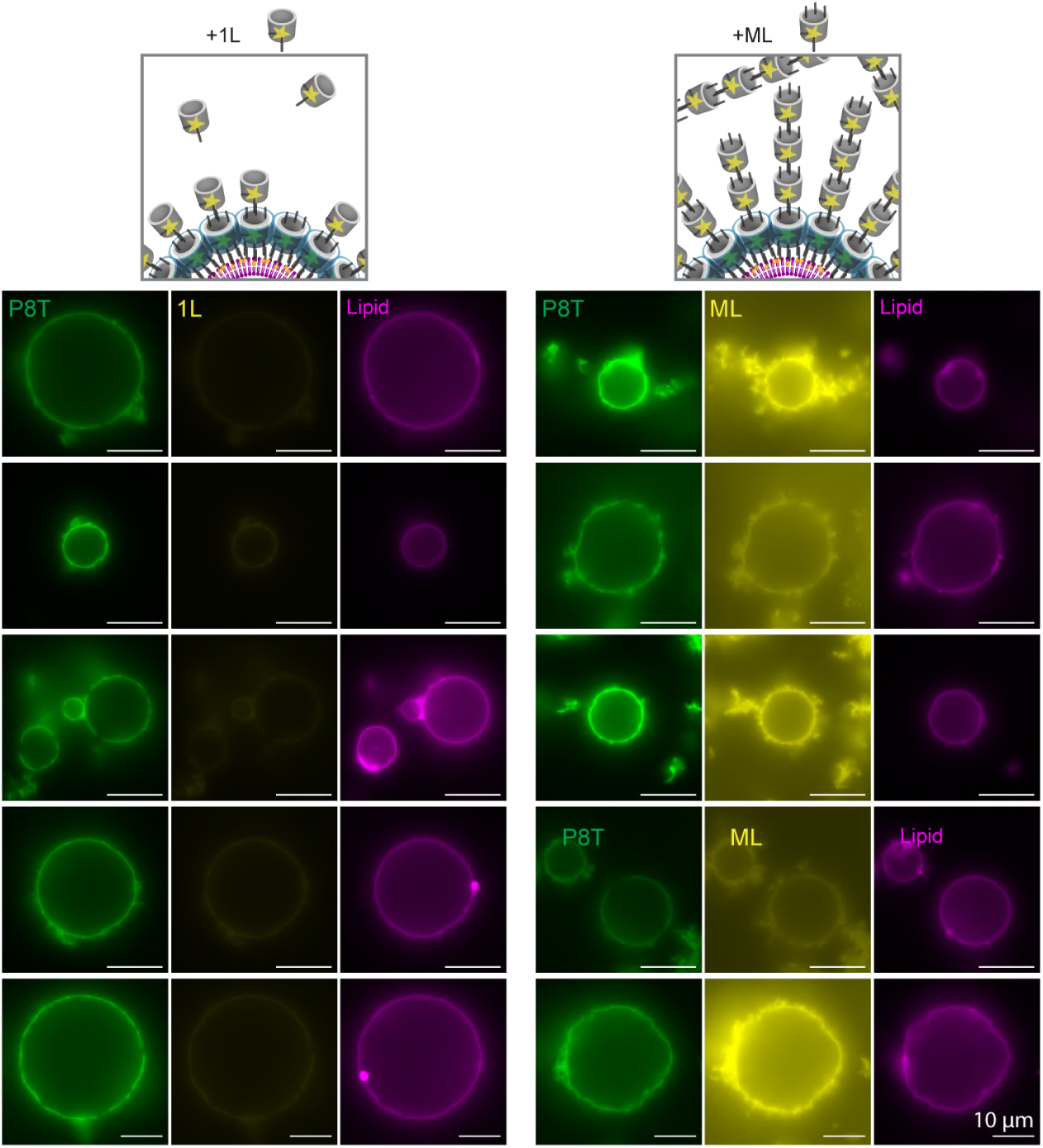
Schematics and corresponding fluorescence microscopy images of P8T Dipids (green), upon reassembly on GUVs (magenta), and subsequent addition of 1L and ML Dipids (yellow) forming bilayer and multilayer DNA shells. Scale bars: 10 µm.

**Figure S10.**
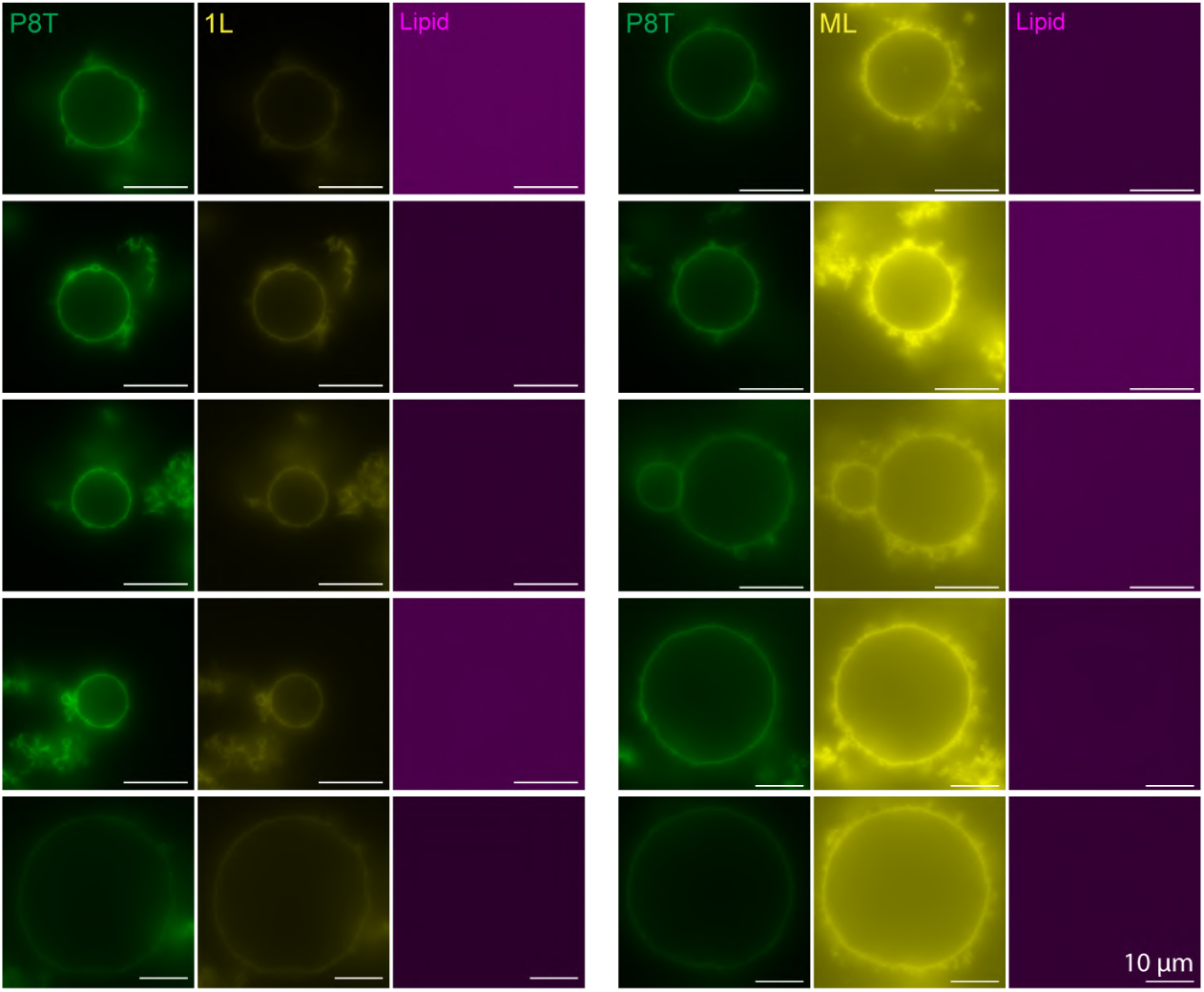
Fluorescence microscopy images of Dipids consisting of P8T (green) and 1L or ML (yellow), upon solubilization of GUVs (magenta) after vesicle-templated assembly, forming freestanding bilayer and multilayer spherical Dipid shells. Scale bars: 10 µm.

**Figure S11.**
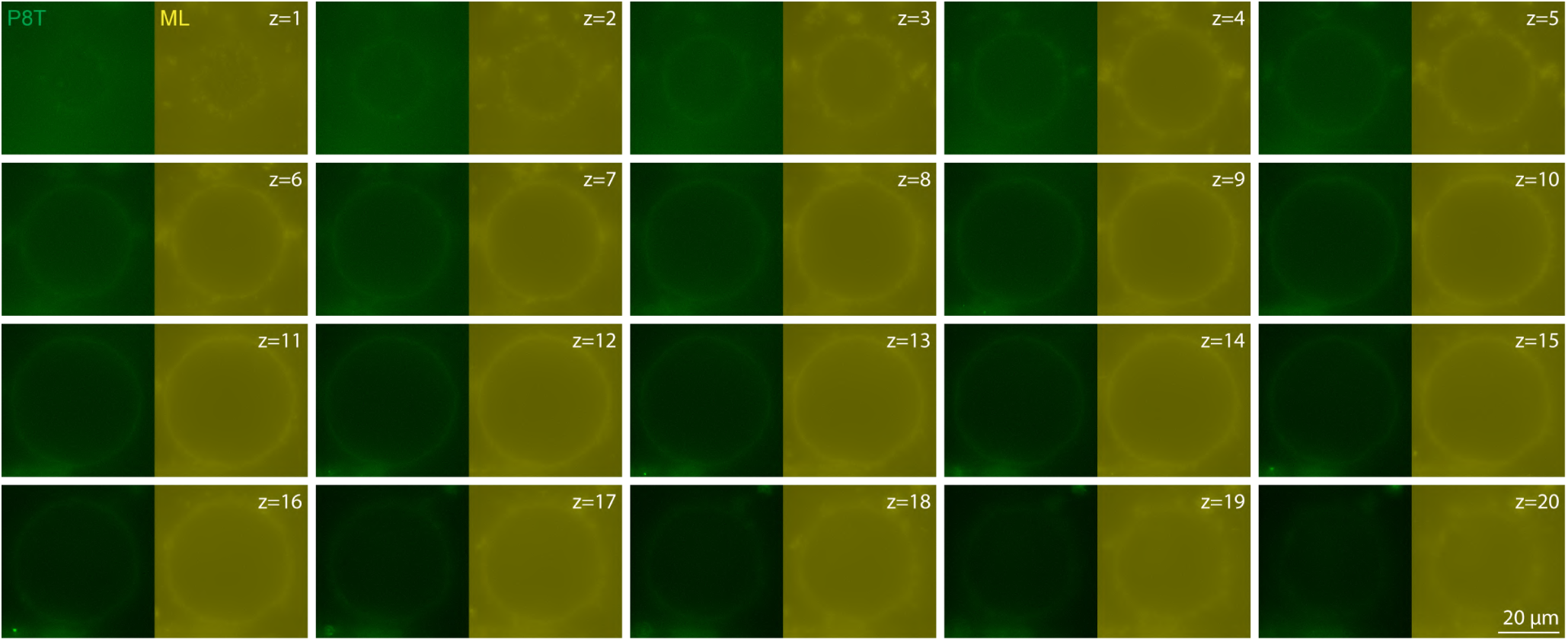
Fluorescence microscopy Z-slices of the freestanding multilayer Dipid shell consisting of P8T (green) and ML (yellow) from Figure 4c (Δ*z* = 2 µm). Scale bar: 20 µm.

**Figure S12.**
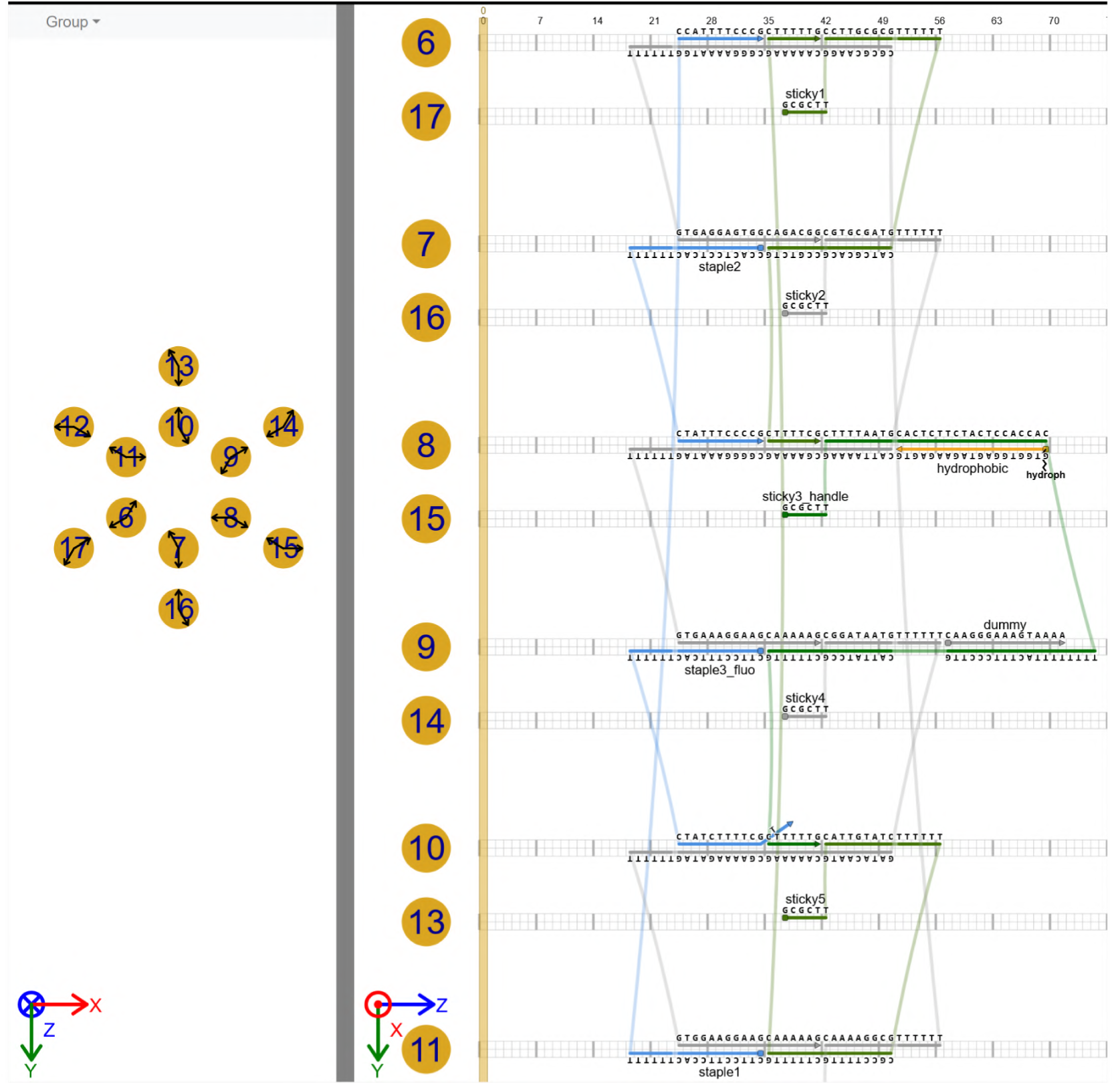
DNA strand routing of the minimal monomer in scadnano [39]. The design file is available on our [GitHub repository](https://github.com/ckarfusehr/Vesicle-templated_self-assembly_of_freestanding_multi-micron_DNA_shells).

**Figure S13.**
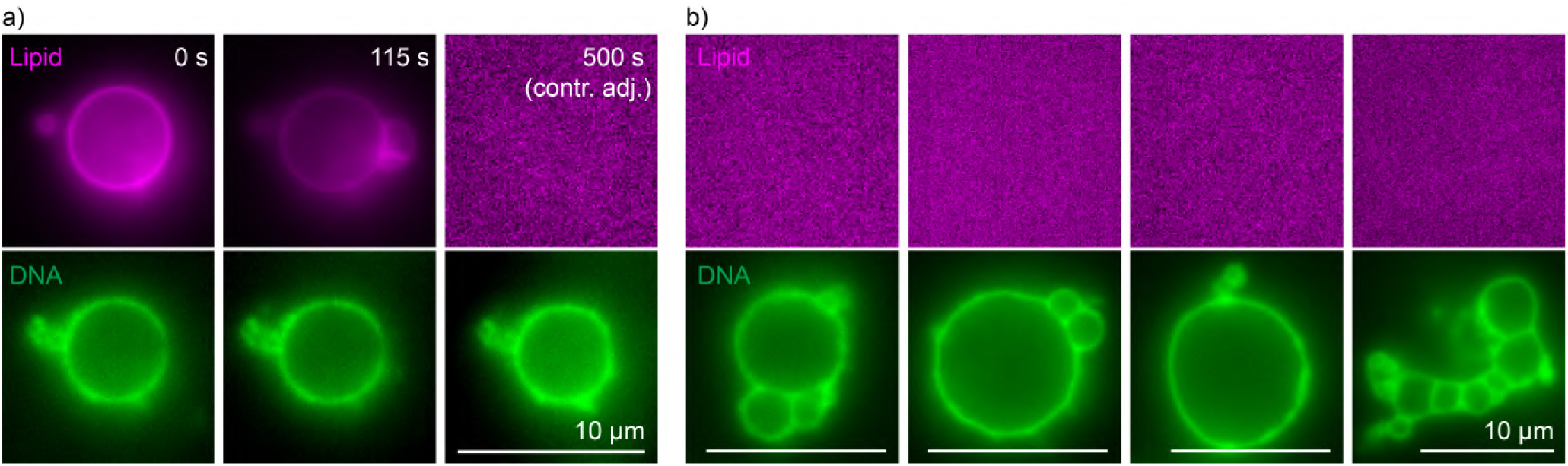
Freestanding DNA shells formed by the minimal monomer. a) Fluorescence time series showing the release of DNA shell (green) from GUVs (magenta). Triton X–100 was added to one side of a narrow channel containing DNA-enclosed GUVs and allowed to diffuse. *t* = 0 s marks the start of acquisition. b) Freestanding DNA shells shown in Figure 4c, next to the lipid signal. Scale bars: 10 µm.

